# Role of the Matrix-Capsid Cleavage Site Polymorphism S124V of HIV-1 Sub-subtype A2 in Gag Polyprotein Processing

**DOI:** 10.1101/2020.11.14.382879

**Authors:** Carla Mavian, Roxana M Coman, Ben M Dunn, Maureen M Goodenow

## Abstract

Subtype C and A HIV-1 strains dominate the epidemic in Africa and Asia, while sub-subtype A2 is found at low frequency only in West Africa. To relate Gag processing *in vitro* with viral fitness, viral protease (PR) enzymatic activity and *in vitro* Gag processing were evaluated. The rate of sub-subtype A2 Gag polyprotein processing, as production of the p24 protein, was reduced compared to subtype B or C independent of PR subtype, indicating that subtype A2 Gag qualitatively differed from other subtypes. Introduction of subtype B matrix-capsid cleavage site in sub-subtype A2 Gag only partially restored the processing rate. Unique amino acid polymorphism V124S at the matrix-capsid cleavage site, together with other polymorphisms at non-cleavage sites, are differentially influencing the processing of Gag polyproteins. This genetic polymorphisms landscape defining HIV-1 sub-subtypes, subtypes and recombinant forms are determinants of viral fitness and frequency in the HIV-1 infected population.

**Graphical Abstract:** 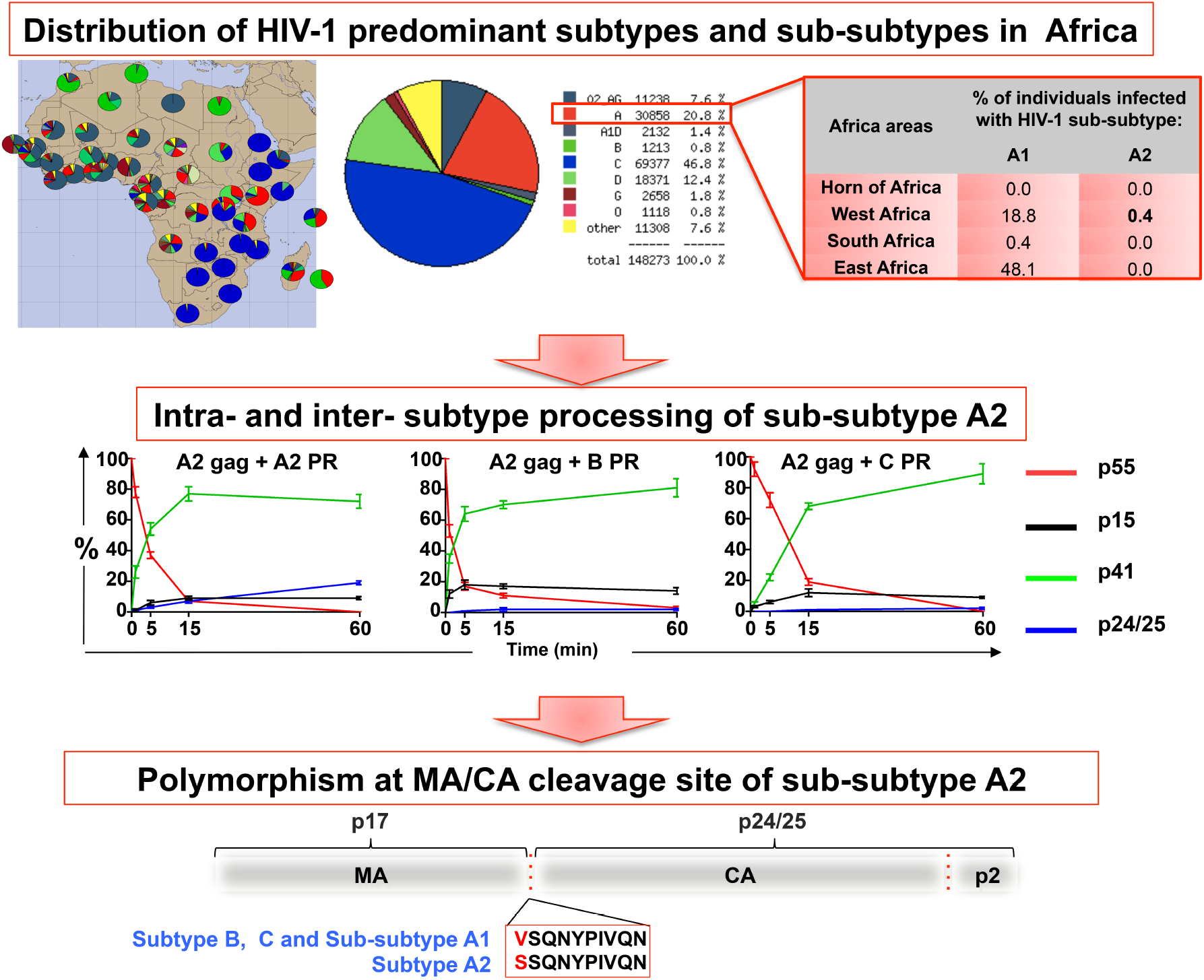

**Highlights:**

1. The polymorphism at matrix-capsid cleavage site, together with non-cleavage sites polymorphisms, direct the processing rate of the substrate, not the intrinsic activity of the enzyme.
2. The less prevalent and less infectious sub-subtype A2 harbors the matrix-capsid cleavage site polymorphism that we report as a limiting factor for gag processing.
3. Sub-subtype A2 Gag polyprotein processing rate is independent of the PR subtype.

## Introduction

High genetic variability and extensive heterogeneity are the major characteristic of HIV-1. It has been recently proposed that the recombinant viruses seeded the early global HIV epidemic in central Africa since the early 1980s [1]. Multiple HIV-1 subtypes co-circulate within regions sharing human interchange/migration, leading to frequent inter-subtype recombination and appearance of circulating recombinant forms (CRFs) (http://www.hiv.lanl.gov/ 2017) [2, 3]. During the last 30 years, 20% of HIV-1 genotypes have undergone inter-subtype recombination and CRFs such as CRF01_AE, CRF_02AG and CRF_07BC contribute to almost 10% of total HIV-1 infections worldwide (http://www.hiv.lanl.gov/ 2017) [2, 4–6]. While global HIV epidemic is characterized by compartmentalized local epidemics dominated by a single subtype indicating strong founder effects, in Africa high diversity of subtypes and CRFs are dominating the epidemic [7–9]. There is no single factor that accounts for geographic prevalence of HIV subtypes observed in countries in Africa; whether high prevalence of certain HIV subtypes reflects increased fitness, or confounding factors, such as sexual transmission networks, ethnicity, socio-economic status or environmental limitations, remains unclear [10, 11]. Mapping recombination sites in the genome of recombinant viruses provides insights into functional regions of the virus genome. For example, recombination between the envelope [*env*] and *gag/pol* regions of viral genomes may increase fitness as 25 to 29% of HIV-1 infected individuals in Africa carry CRF with discordant *gag* and *env* subtypes, predominantly subtypes A and G [4, 5]. In contrast, recombination within *gag* and *pol* regions between subtypes may have a fitness cost to the virus, as the *gag-pol* region coevolves as a functional unit reflecting the interplay between the enzymatic activity of PR in *pol* and the cleavage-site substrates distributed across the Gag polyprotein [12, 13].

The evolution of HIV-1 subtype A, similarly to subtype F, gave rise to five different distinct lineages, defined as sub-subtypes [A1, A2, A3, A4, A5] [14–17]. These sub-subtypes are highly related to the parental subtype A clade, and form sub-clades with a distinct sister clade to subtype A in phylogenetic trees from *gag*, *pol*, *env*, and *nef* regions with genetic distance of about half of that between subtypes [14]. Sub-subtype A2, the second most prevalent sub-subtype of the A clade, was originally described in Kenya and sporadically reported in other parts of the world [14, 18–20]. In contrast to sub-subtype A1, prototypic for the subtype A epidemic worldwide according to the new nomenclature, sub-subtype A2 is found to infect 0.4% of African population only in West Africa [14, 21].

The extensive HIV-1 subtype diversity found in Africa, epicenter of the pandemic, offers the best frame to relate genetics of subtypes to socio-behavioral factors influencing viral fitness [2, 3]. Infections by non-B subtypes of HIV-1, such as subtypes C and A, predominate in specific regions in Africa [2]. However, because subtype B dominated western hemisphere countries HIV-1 infections, antiretroviral drug development and susceptibility testing were originally targeted to subtype B, and consequently molecular studies on non-B subtypes Gag polyproteins and PR processing are lacking [22–27]. Viral fitness research aimed to identify polymorphisms that play a key role in antiretroviral mechanisms, have focused mainly on replicative fitness within hosts or in cultured cells [28]. Assessing viral fitness is complex to dissect at the whole virus level, as compensation across the genome can provide an “apparently” fit virus, even though certain functional aspects of the virus may be suboptimal. We and others have shown that natural genetic polymorphisms present at Gag cleavage sites can modulate Gag processing and relate to fitness in different ecosystems [i.e., drug resistant or drug sensitive virus in the absence or presence of PI] [13, 22, 25, 29, 30]. The idea underlying our approach is the characterization of the role of Gag polymorphisms *ex vivo,* which we expect to be correlated with fitness, as *ex vivo* provides insights into potential functional differences that may be obscured by replication competent *in vitro* assays.

Our study investigates the role of polymorphisms found at the cleavage site of sub-subtype A2, B and C Gag polyproteins, during *ex vivo* processing by the viral protease, rather than to viral replicative analysis *in vivo*, and relates the Gag processing events *ex vivo* with viral fitness in human populations and geographic prevalence [22, 25].

## Materials and methods

### Mutagenesis and expression of HIV-1 PR and *gag-pol* genes

HIV-1 subtype B PR allele was obtained from a molecular clone of HIV-1AD [13, 31], while sub-subtype A2 (NIH clone p92UG037.1, accession number AF286237) and subtype C alleles (NIH clone p94IN476.104, accession number AF286223) were obtained through the AIDS Research and Reference Reagent program, Division of AIDS, NIAID, from Drs. Rodenburg, Gao, and Hahn [14, 32]. HIV-1 PR cloning into the pET23a expression vector (Novagen), expression in *Escherichia coli* strain BL21 Star DE3 PlysS (Invitrogen), and purification from inclusion bodies were performed as previously described [33, 34].

The *gag-pol* genes from subtypes B, C, or A2 were amplified using primers engineered to introduce restriction sites XhoI at the 5’ end and MluI at the 3’ end of the amplicon for directional cloning into the TNT expression vector. Sequence alignment of PR and Gag of sub-subtype A2, subtypes B or C, and of subtype A1 (Genbank accession number: AB098332), was performed with Clustalw2 (http://www.ebi.ac.uk/Tools/msa/clustalw2/) (Figure S1).

The subtype B matrix/capsid (MA/CA) cleavage site was introduced into sub-subtype A2 p55 *gag-pol* gene either by changing residue 124 from valine (V) to serine (S) (A2gagS124V) or introducing the glutamine-valine (QV) dipeptide at position 124 (A2gagQV). Mimicking the sub-subtype A2 MA/CA cleavage site in the subtype B Gag polyprotein was performed by mutating the valine in position128 to either leucine (L) (BgagV184L) or S (BgagV128S), or by deletion of the glutamine and valine residues in positions 127 and 128 (Bgag_ΔQV). Mutations were confirmed by Sanger sequencing at the Interdisciplinary Center for Biotechnology Research of the University of Florida.

### PR activity constants

The Michaelis-Menten constants k_cat_, K_m_, and k_cat_/K_m_ values were determined *in vitro* for each PR subtype variant using the chromogenic substrates K-A-R-V-nL*Nph-E-A-nL-G, which resembles the CA/p2 cleavage site of subtype B [35], as previously described [33]. Cleavage of the substrate was monitored using a Cary 50 Bio Varian spectrophotometer equipped with an 18-cell multi transport system.

### PR processing in trans of Gag polyproteins transcribed and translated *in vitro*

TNT plasmids containing a *gag* open reading reproducing exactly the encoded Gag polyproteins of subtypes B and C, and sub-subtype A2 were mixed with TNT T7 Quick Master Mix (rabbit reticulocite lysate and a mixture of all the amino acids except methionine), [^35^S] Met (1000 Ci/mmol at 10 mCi/ml) and nuclease-free H_2_O. After incubation at 30°C for 2 h, active HIV-1 PR at concentrations ranging from 10 nM to 50 nM was added to the reaction mixture. Aliquots were collected immediately before [time 0] and at 1, 5, 15, 60 and after adding enzyme, quenched with 2x Laemmli buffer at 1:1 ratio and heated to 70°C for 2 min. An extra aliquot was sampled at 90 min for recombinant Gag proteins. Samples were electrophoresed through a 10-20% SDS-PAGE gel (BioRad) that was subsequently fixed for 30 min in a solution containing 10% acetic acid and 5% hydrochloric acid and then soaked for 5 min in 10% glycerol solution. Gels were dried and exposed to either a XAR-5 film (Kodak) or a phospho-screen at room temperature. The amount of labeled proteins was quantified using a Molecular Dynamics PhosphorImager, Storm 860 model and Image Quant (Promega). Total intensity of bands of interest was considered 100 and the amount of each product was calculated based on number of Met residues present in each subtype Gag polyprotein as percentage of the total amount of labeled substrate in the lane. Rate of processing was compared across subtypes and experiments as total production of the p24 protein at 60 min.

## Results

### Kinetic analysis of HIV-1 PR alleles from sub-subtype A2, subtypes B and C

Polymorphisms found within the HIV-1 PR reflect differences between and within subtypes sub-subtype A2, subtype B, and subtype C. Polymorphic residues found among the HIV-1 PR subtypes were residue 37 within the elbow region (aa 37-44), residue 69 in the 60s’ loop (aa 66-69), and residues 15 and 16 wihtin the 10s’ loop (aa15-18) (Supplementary Figure 1A). No Polymorphic residue was located within the active site or the dimerization region of the PR (Supplementary Figure 1A). Efficiency of processing by each PR was assayed using a substrate that mimics the conserved CA/p2 cleavage site of the Gag polyproteins [33] (Supplementary Figure 1B). Subtype B PR displayed the highest k_cat_ values, subtype A2 PR had the highest K_m_, while subtype C PR displayed the lowest values for k_cat_ or K_m_ (Table 1). Overall, the catalytic efficiency (k_cat_/K_m_) of subtype A2 or subtype C PR was about 74% or 62%, respectively, of the activity of subtype B PR (Table 1).

**Table 1.** Kinetic parameters of HIV-1 PRs of subtypes B, C and sub-subtype A2.

| **Subtype** | **$K_m$ ($\mu\text{M}$)** | **$k_{\text{cat}}$ ($\text{s}^{-1}$)** | **$k_{\text{cat}}/K_m$ ($\text{s}^{-1} \mu\text{M}^{-1}$)** |
| --- | --- | --- | --- |
| **B** | $19.5 \pm 2$ | $10 \pm 1$ | $0.51 \pm 0.06$ |
| **C** | $17 \pm 1$ | $5.6 \pm 0.2$ | $0.32 \pm 0.02$ |
| **A2** | $22 \pm 2$ | $8.4 \pm 0.4$ | $0.38 \pm 0.04$ |

Standard deviations are best-fit values through multiple points done twice.

### HIV-1 Gag polyprotein processing is independent of PR subtype

The cleavage sites for capsid (CA) and p2 [CA/p2], and nucleocapsid (NC) and p1 [NC/p1] were conserved among the three subtypes, while the cleavage site for p2 and NC [p2/NC] was polymorphic in each subtype. Two cleavage sites where identical in B and C Gag subtypes, but different from A2 Gag, and these were the cleavage sites for matrix/capsid [MA/CA] and for p1/p6Gag (Supplementary Figure 1B). Particularly, the polymorphic residue in the MA/CA cleavage site was found at the first residue of the site: a valine residue in B and C Gag subtypes (aa V128) and a serine residue in A2 Gag (S124) (Supplementary Figure 1B).

To assess the contribution of polymorphisms found in Gag polyproteins to Gag processing, we performed an intra-subtype Gag processing of sub-subtype A2, subtypes B and C Gag polyproteins by their corresponding PR subtype, taking in account the accumulation of p24 as indicator of processing rate (Figure 1). Subtype B p55 was processed with an approximate half-life time (T_1/2_) of 1 min (Figure 1, Supplementary Figure 2A-C). Approximately 50% production of p41 was reached at 2 min and peak production of p41 by 15 min. Accumulation of Subtype B p24 reached approximately 25% by 60 min. Subtypes C p55 was processed slower (T_1/2_ of 4 min) as compared to subtype B, and 50% production of p41 was reached at 5 min, and peak production of p41 by 15 min. However, as respect to subtype B, production of subtype C p24 was faster than subtype B, with approximately 60% p24 accumulated by 60 min. Sub-subtype A2 p55 decline (T_1/2_ of 4 min), and 50% p41 production (4 min) and peak (15 min) were similar to subtype C. However, sub-subtype A2 final production of p24 was slower as compared to subtypes B and C, with less than 20% accumulation by 60 min (Figure 1, Supplementary Figure 2A-C).

**Figure 1.**
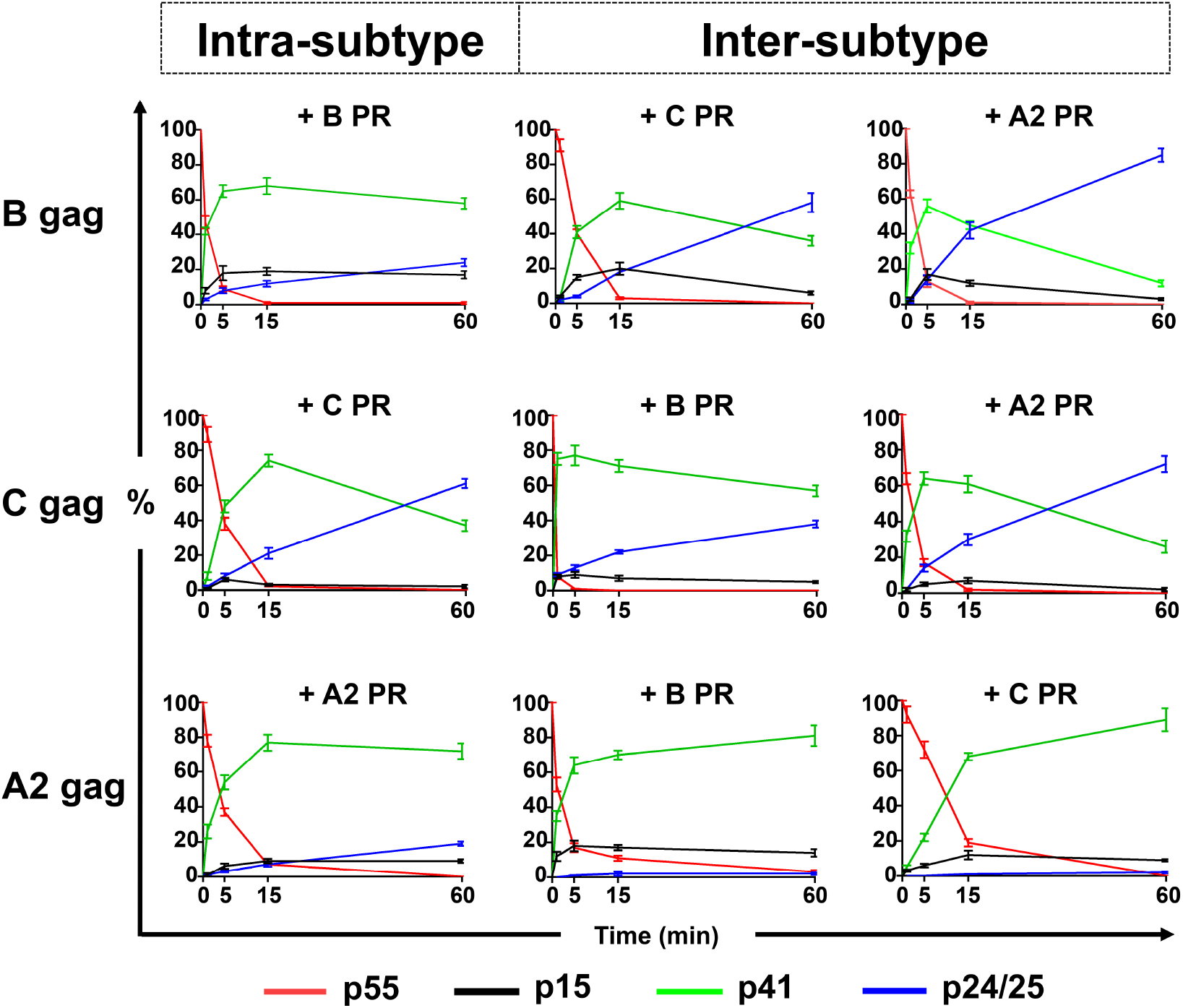
*In trans* processing of HIV-1 subtype B, C and sub-subtype A2 Gag polyproteins by same and different PR subtype. Graphs show the percentage of substrate and products derived from the *in trans* processing of Gag polyprotein subtype B, subtype C and sub-subtype A2 upon addition of intra-subtype and inter-subtype PR combinations. Data are based on at least 3 experiments and expressed as mean ± SEM. Representative gel experiment is reported in Supplementary Figure 2A-C.

Comparing the inter-subtype processing of subtype B Gag by either sub-subtype A2 or subtype C PRs, the decline of p55 was slower with T_1/2_ of 3 or 4 min, respectively. The production of p41 was slower as well, with 50% production reached around 5 min with both PRs. However, overall production of subtype B p24 was increased, with 60% production by sub-subtype C PR and over 80% by subtype A2 PR at 60 min. Subtype C Gag showed faster kinetics when processed by subtype A2 or subtype B PRs, with T_1/2_ p55 decline of 1 min. Peak subtype C p41 production by subtype B PR was of 1 min, and of 5 min when processed by sub-subtype A2 PR, as by subtype C PR. Production of p24 by sub-subtype A2 PR was increased by 10% at 60 min as compared to subtype C PR processing amount, whereas decreased by 20% when processed by subtype B PR. The increased p41 production over the course of the experiment, and especially as shown for sub-subtype A2 p41 production by subtype C PR, indicates that the PRs were active during 60 min.

Rate of sub-subtype A2 p55 processing was slower when processed by subtype C PR (T_1/2_ of 8 min), and faster when processed by subtype B PR (T_1/2_ of 2 min) (Figure 1, Supplementary Figure 2A-C). Rate of production of sub-subtype A2 p41 was increased by subtype B PR processing, but not by subtype C PR. However, sub-subtype A2 Gag processing by subtypes B or C PRs decreased the accumulation of sub-subtype A2 p24 at 60 min to 2%. Overall, the amount of p24/25 sub-subtype A2 Gag processing resulted from processing by subtype A2 PR (intra-subtype) was 2-to 5-fold less as compared to the amounts generated by respective intra-subtype processing of subtypes B or C Gags. The rate of inter-subtype sub-subtype A2 Gag processing was 15-to 20-fold lower if compared to the inter-subtype processing of subtypes B or C. Moreover, the amount of p24/25 produced by the intra-processing of sub-subtype A2 Gag was 4-fold less than the amount resulted from inter-subtype of subtype B Gag by sub-subtype A2 PR. This latter observation suggests that sub-subtype A2 PR activity was reduced in presence of sub-subtype A2 Gag polyprotein but efficient in presence of other Gag subtypes (Figure 1, Supplementary Figure 2A-C).

Together these results indicate that sub-subtype A2 Gag p55 can be processed by subtype B PR even more rapidly than by A PR, perhaps reflecting the modest polymorphisms in p2/NC cleavage site; and sub-subtype A2 Gag p41 accumulates even more rapidly when processed by B PR than by A PR (Supplementary Figure S1A). Finally, independent of which subtype PR. sub-subtype A2 p24/p25 was not accumulated at the same rate as for subtypes B or C products, and a possible explanation is the single amino acid polymorphism at the MA/CA cleavage site (Supplementary Figure S1A).

### The V124S polymorphism at the MA/CA cleavage site of sub-subtype A2 Gag polyprotein is influencing Gag processing rate

The MA/CA cleavage site harbored the V124S polymorphism on residue S124 of sub-subtype A2 Gag polyprotein, corresponding to residues V128 or V125 of subtype B or C, respectively (Figure 2A, Supplementary Figure 2D). To assess the role for V128/S124 polymorphism in determining p55 processing rate, the subtype B or sub-subtype A2 MA/CA cleavage sites were mutated into sub-subtype A2 p55 (A2gagS124V and A2gag_QV) or subtype B Gag polyprotein (BgagV128L, BgagV128S and Bgag_ΔQV) respectively, and all mutant proteins were processed by subtype B PR (Figure 2B, Supplementary Figure 2E). The rate of processing of the A2gagS124V and Agag_QV mutant proteins by subtype B PR increased by two-fold, with twice as much p24/25 production when compared to wild type, but seven fold less compared to subtype B Gag. On the other hand, the processing of the subtype B p55 mutants decreased significantly: p24/25 production by processing the BgagV128L mutant protein was double as compared to wild-type, while by processing the BgagV128S mutant was three times less when compared to wild-type subtype B Gag, and twice as much as for sub-subtype A2 Gag. The amount of p24/25 generated processing Bgag_ΔQV and BgagV128S mutant proteins was similar. The constant increased p41 production indicates that the PRs were active at least for 90 min.

**Figure 2.**
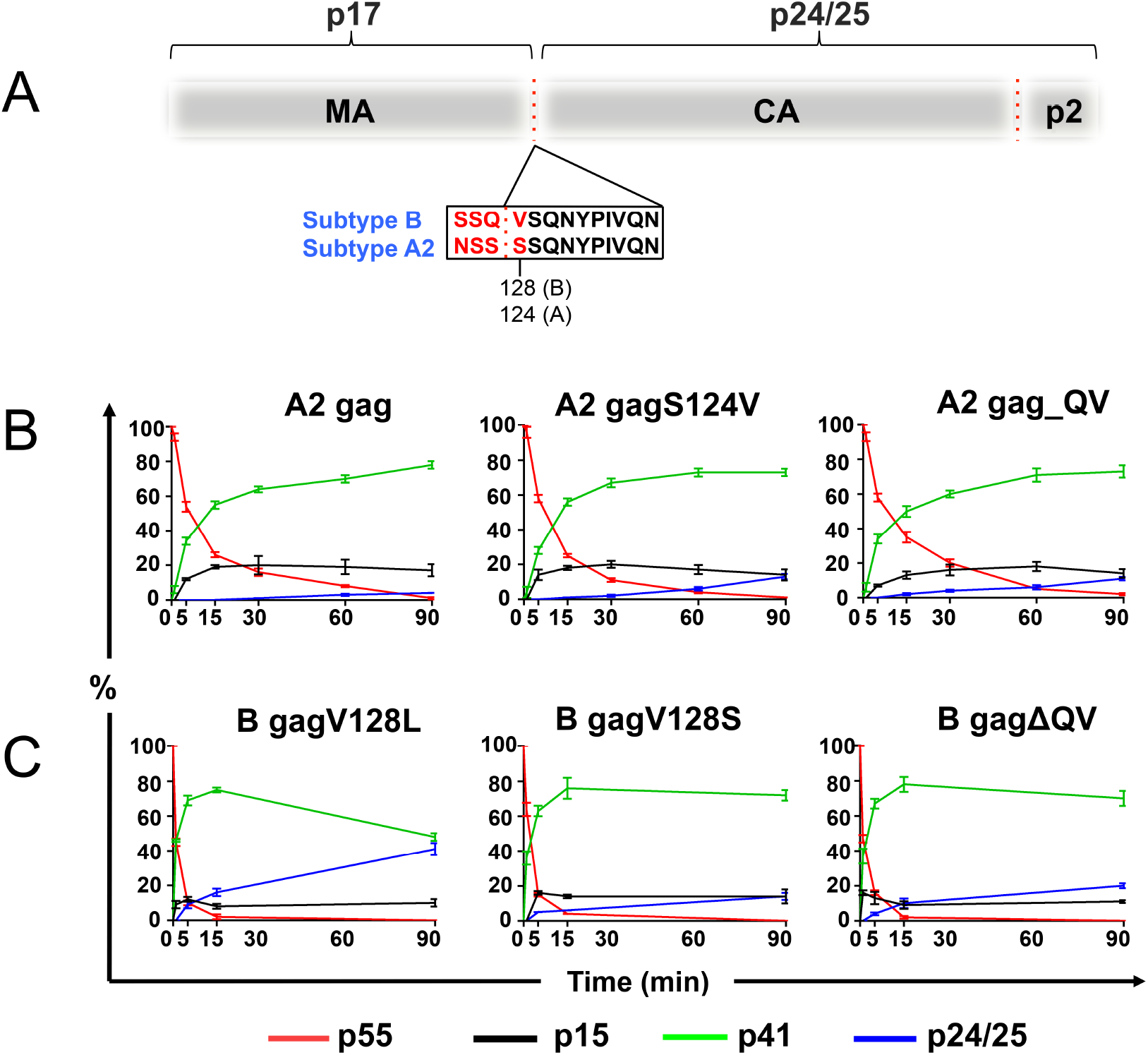
*In trans* processing of HIV-1 sub-subtype A2 and subtype B gag polyprotein mutants by subtype B PR. (A) Schematic representation of the p55 gag precursor with cleavage site for matrix (MA) and capsid (CA) [MA/CA], cleavage site for production of p24/25. The MA/CA cleavage site sequence for subtype B and sub-subtype A2 are shown in the box, conserved residues are reported in black and polymorphic residues in red. (B) Graphs show the percentage of substrate and products derived from the processing of Agag, AgagS124V, AgagQV polyproteins, and BgagV128L, BgagV128S, BgagΔQV polyproteins upon addition *in trans* of subtype B PR is shown in time. Data are based on at least 3 experiments and expressed as mean ± SEM.

Taken together, these results indicated that the V124S polymorphism at the MA/CA cleavage site affected the processing rate of sub-subtype A2 Gag polyprotein, and suggested that other polymorphisms within Gag may also contribute to regulate processing. Overall these results confirmed that the MA/CA cleavage site plays a role in regulating HIV-1 Gag polyprotein processing rate.

## Discussion

The global distribution of HIV-1 is a dynamic process determined by viral genetic diversity due to high mutation and recombination rate. Viral fitness of dominant subtypes and regional shifting in distribution of non-B subtypes and recombinants is occurring, especially in regions of Sub-Saharan Africa and Southeast Asia. Sub-subtype A2, that found its niche in West Africa in 2000 and is currently infecting 0.4% of the West African population, has been almost entirely replaced by the more infectious sub-subtype A [21, 36]. The geographical constriction of A2 to local epidemics and the predominance of sub-subtype A1 over A2, as well as other subtypes, may be due to transmission bottlenecks reflecting viral genetic effects as well as social/behavioral or environmental limitations.

The goal of our study was to relate the Gag processing events *ex vivo* with viral fitness in human populations, rather than to viral replicative analysis *in vivo*. Sub-subtype A2 harbors a serine residue in position 124 which we described as a limiting factor of Gag processing; the sub-subtype A1 harbors instead a valine residue, as subtypes B and C [14]. Mutations at the MA/CA cleavage site are reported to reduce viral infectivity explaining the high degree of conservation of the MA/CA cleavage site residues within group M subtypes and recombinants [22, 37]. While a 98% of conservation is found among subtype A, A1, B and C, sub-subtype A2 showed only 89% [22].

CRFs are emerging in the HIV-1 epidemic representing the new direction of HIV-1 evolution. Among the diversity of CRFs worldwide, in Africa only two CRFs show exchange between the subtype A gag polyprotein and subtype C PR and vice versa as result of the recombination events [38]. A third CRF was found in Canada harboring subtype C gag polyprotein and subtype A PR [39].

By combining Gag polyproteins and PRs from different subtypes, our study examines the natural subtype-interplay processing event found during infection with CRFs that carry PR and Gag polyproteins from different HIV-1 subtypes. Our finding showed that non-A2 subtype PR processes the Gag polyproteins of subtype A2 with a higher rate as compared with its own PR. CRFs that harbor the *gag* region of subtype A present a tendency not to conserve the corresponding *pol* region and vice versa, which harbors the PR of subtype A but the gag polyprotein of subtype K [22]. Given our results, it is not surprising that two recombinant forms of sub-subtype A2 with subtype D found in Kenya, CRF16_A2D and CRF21_A2D, are preserving the MA/CA cleavage of sub-subtype A2 (1186 nt) and encoding the subtype D PR (2253-2550 nt) (http://www.hiv.lanl.gov/ 2017) [38, 40]. The CRFs of sub-subtype A2 and subtype D, CRF16_A2D and CRF21_A2D, prove recombination as alternative mechanism of spread within the population for strains such as sub-subtype A2 which otherwise would be relatively rare in the pandemic. Moreover, the incidence of the CRFs of sub-subtype A2 with subtype D (2-9%) among HIV-1 infections in Kenya, lower as compared to subtype D (28%) and higher to sub-subtype A2 (2.7%) [41], is in agreement with our hypothesis that sub-subtype A2 MA/CA cleavage may reduce viral fitness. Together with our findings, these data suggest that compensation of subtype D PR for the sub-subtype A2 MA/CA cleavage site is insufficient to improve the processing rate. Monitoring of sub-subtype A2, A2-containing recombinants, as well as other rare subtypes circulating in limited geographic area, together with consideration of the socio-economic factors that influence transmission of HIV-1, may provide fundamental information for further development and evaluation of candidate niche vaccines to eradicate even minor variants.

In conclusion, our findings suggest that the V124S polymorphism present in the MA/CA cleavage of sub-subtype A2, together with the amino acidic context surrounding the cleavage and at non-cleavage sites, is a novel key factor influencing sub-subtype A2 fitness and a potential mechanisms governing PR interactions with Gag, which substantially influence the frequencies of HIV-1 subtypes, sub-subtypes and CRFs worldwide.

## Conclusions

- The processing rate of Gag polyproteins of sub-subtype A2, as well as of subtypes B and C, is independent of the intrinsic activity of the PR or subtype.
- The S124V polymorphism present at the matrix-capsid cleavage site of sub-subtype A2 influence the direct processing rate of the Gag polyprotein together with non-cleavage sites polymorphisms.
- The less prevalent and less infectious HIV-1 sub-subtype A2 harbors a matrix-capsid cleavage site polymorphism that is a limiting factor for gag processing, while the more prevalent and infectious HIV-1 sub-subtype A1 present the same matrix-capsid cleavage site as subtypes B and C.
- The analysis on sub-subtype A2, and subtypes B and C, that we performed suggested that the viral fitness found *in vivo* among from different subtypes in the “wild” among HIV-1 infected individuals may be related to a lower Gag processing rate.

## Supplementary Material

**Table S1.** Geographical prevalence and distribution of HIV-1 sub-subtypes A1 and A2.

| Geographic areas | Individuals infected with HIV-1 sub-subtype (%) |  |
| --- | --- | --- |
|  | A1 | A2 |
| Horn of Africa | 0.0 | 0.0 |
| West Africa | 18.8 | 0.4 |
| South Africa | 0.4 | 0.0 |
| East Africa | 48.1 | 0.0 |
| North America | 1.0 | 0.0 |
| South America | 0.1 | 0.0 |
| North Asia | 99.1 | 0.0 |
| Central Asia | 0.0 | 0.0 |
| South Asia | 0.0 | 0.0 |
| Europe | 4.7 | 0.0 |

<sup>a</sup>Information retrieved from [21]

**Figure S1.**
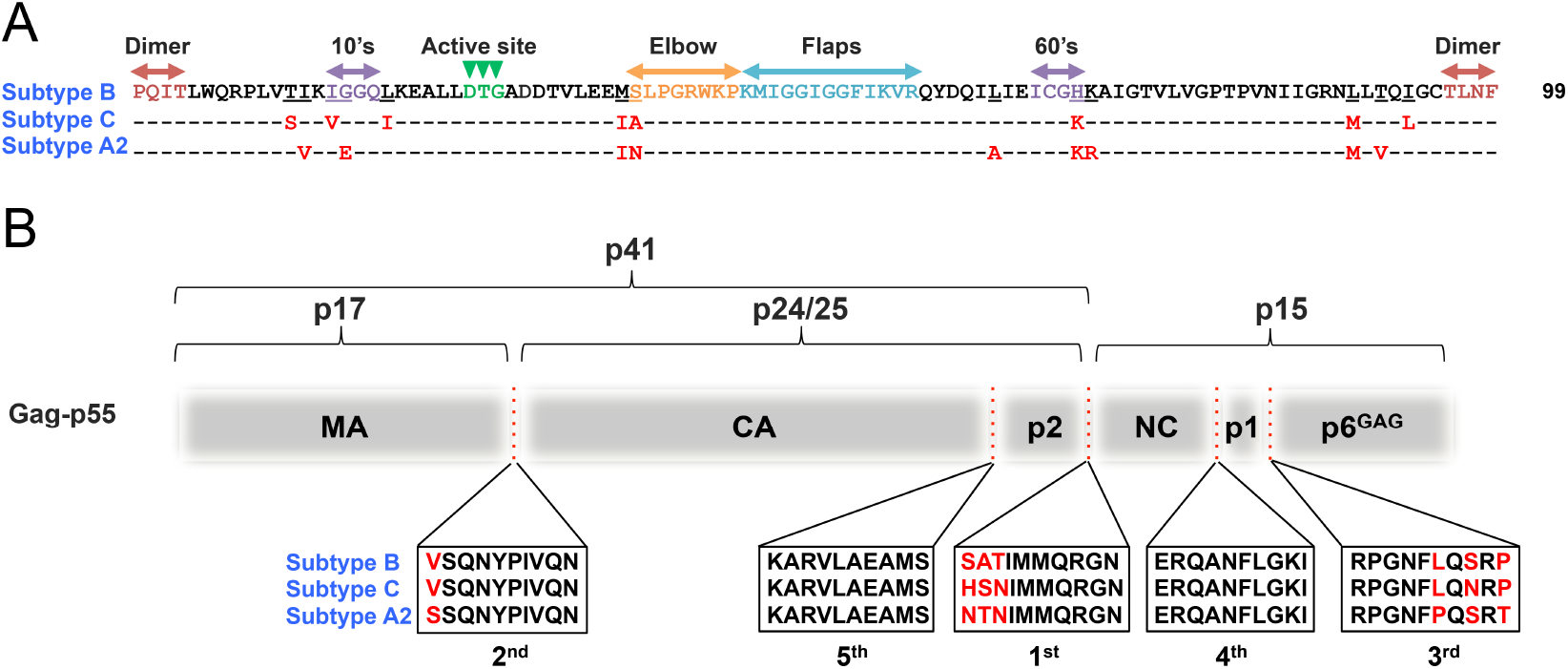
Schematic representation and sequence alignment of subtype B, C and sub-subtype A2 HIV-1 PR and Gag polyproteins with polymorphisms highlighted. (A) Sequence alignment for subtype B, C and sub-subtype A2 PRs is reported compared to subtype B sequence as reference. Same amino acid residues are shown with -,polymorphisms in red, and the catalytic residue in green. (B) Schematic representation of subtype B, C and sub-subtype A2 Gag polyproteins with cleavage sites for matrix (MA), capsid (CA), p2, nucleocapsid (NC), p1 and p6^GAG^ proteins sequences reported in boxes. Numbers report the order of Gag processing. Conserved residues are shown in black, polymorphic residues in red.

**Figure S2.**
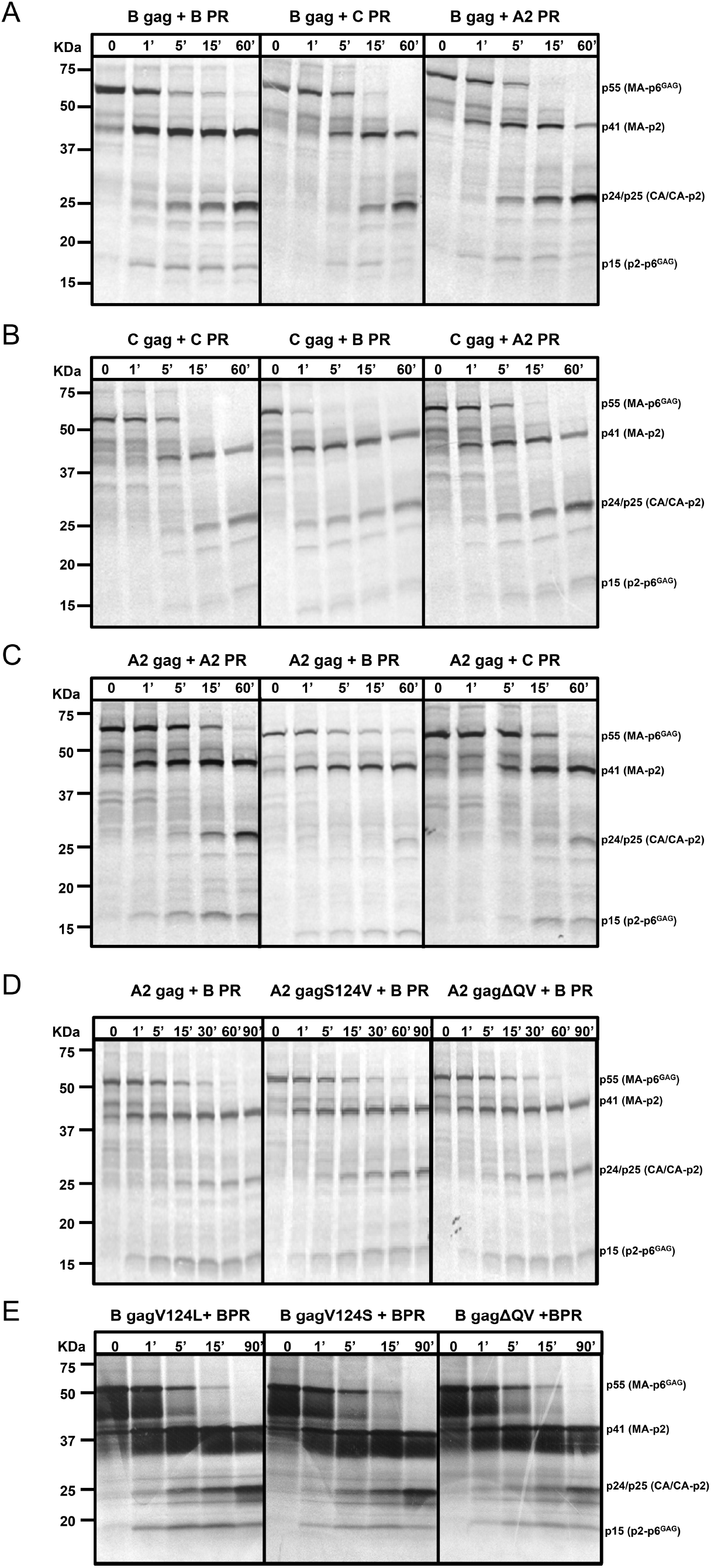
Gels for *in trans* processing of HIV-1 subtype B, C and sub-subtype A2 Gag polyproteins by same and different PR subtype, and for *in trans* processing of HIV-1 subtype B and sub-subtype A2 gag polyprotein mutants by subtype B PR. (A) Processing of Gag polyprotein subtype B, (B) subtype C and (C) sub-subtype A2 upon *in trans* addition of different subtype PR. (D) processing of gag polyprotein sub-subtype A2, and of AgagS124V, AgagQV, (E) BgagV128L, BgagV128S, and BgagΔQV mutant gag polyproteins upon addition *in trans* of subtype B PR. Cocktail of gag polyprotein and subtype PR is shown above gel panels, and percentage of substrate and products is reported below. Time is indicated in minutes in wells above corresponding lane.

